# Interpretable high-resolution dimension reduction of spatial transcriptomics data by SpaHDmap

**DOI:** 10.1101/2024.09.12.612666

**Authors:** Junjie Tang, Zihao Chen, Kun Qian, Siyuan Huang, Yang He, Shenyi Yin, Xinyu He, Buqing Ye, Yan Zhuang, Hongxue Meng, Jianzhong Jeff Xi, Ruibin Xi

## Abstract

Spatial transcriptomics (ST) technologies have revolutionized tissue architecture studies by capturing gene expression with spatial context. However, high-dimensional ST data often have limited spatial resolution and exhibit considerable noise and sparsity, posing significant challenges in deciphering subtle spatial structures and underlying biological activities. Here, we introduce SpaHDmap, an interpretable dimension reduction framework that enhances spatial resolution by integrating ST gene expression with high-resolution histology images. SpaHDmap incorporates non-negative matrix factorization into a multimodal fusion encoder-decoder architecture, enabling the identification of interpretable, high-resolution embeddings. Furthermore, SpaHDmap can simultaneously analyze multiple samples and is compatible with various types of histology images. Extensive evaluations on synthetic and real ST datasets from various technologies and tissue types demonstrate that SpaHDmap can effectively produce highly interpretable, high-resolution embeddings, and detects refined spatial structures. SpaHDmap represents a powerful approach for integrating ST data and histology images, offering deeper insights into complex tissue structures and functions.

## Introduction

The revolutionary spatial transcriptomics (ST) technologies have enabled the simultaneous capture of gene expression profiles and spatial locations of cells or spots within tissue samples^1-6^. These cutting-edge techniques have been widely applied across various fields to uncover complex tissue structures, organizations, and microenvironments^7-10^. However, the spatial resolution of commonly used platforms like 10X Visium remains limited. Each ST spot typically contains multiple cells, and large areas of tissue are not covered by sequencing spots. Newer technologies, such as Stereo-seq^2^ and Xenium^3^, achieve much higher resolution but produce much sparser and noisier data. Aggregating spots into larger meta-spots is often needed to increase the signal-to-noise ratio, which unfortunately significantly reduces spatial resolution^11^.

ST data are high-dimensional and often exhibit considerable noise and sparsity, necessitating dimension reduction as a crucial step in their analysis. Since traditional dimension reduction methods do not account for spatial information, specialized methods tailored for ST data have recently been developed^12-17^. By leveraging spatial information, these methods generate spatially-aware, low-dimensional representations that better preserve spatial relationships between cells or spots. Several of these methods, such as NSFH^15^ and SpiceMix^16^, are based on Non-negative Matrix Factorization (NMF) and thus can provide more interpretable, parts-based representations of the high dimensional ST data. However, these available dimension reduction techniques for ST data are constrained by the resolution limitation, facing significant challenges in deciphering subtle expression patterns and complex spatial architectures. Further, when ST data contain multiple samples^18-20^, spots between samples may not have a spatial relationship. Methods that heavily rely on spatial information are not directly applicable to these multi-sample datasets.

ST data are often accompanied by high-resolution histology images, such as Hematoxylin-Eosin (H&E) stained images or immunofluorescence images, which provide valuable information about tissue morphology, cellular composition, and spatial organization at a higher resolution. Several methods that integrate ST gene expression and histology image patches around sequenced spots have been developed for dimension reduction and spatial domain detection^12, 21, 22^. To enhance spatial resolution in ST data, novel computational methods have been developed for high-resolution gene expression recovery and improved tissue architecture identification^23-28^. Nevertheless, these resolution-enhanced methods are designed to address gene expression recovery rather than dimension reduction at high resolution. Furthermore, they primarily focus on integrating H&E images, lacking support for other types of histology images, and do not support simultaneous analysis of multiple samples.

In this paper, we present SpaHDmap (deep fusion of <u>spa</u>tial transcriptomics and histology images for interpretable <u>h</u>igh-<u>d</u>efinition embedding <u>map</u>ping), a multi-modal dimension reduction framework to generate interpretable, high-resolution embeddings of ST data by leveraging histology images. SpaHDmap embeds the NMF of spatial gene expression into a multi-modal encoder-decoder architecture and learns high-resolution embeddings through the accurate recovery of both spatial gene expression and histology images. This versatile framework can seamlessly handle multiple samples simultaneously and support various types of histology images. We benchmarked SpaHDmap using 2 synthetic datasets and 12 real ST datasets from various technologies and tissue types, including 5 mouse brain datasets, 4 medulloblastoma datasets from patient-derived orthotopic xenograft (PDOX) mice, and 3 newly sequenced human colorectal cancer (CRC) datasets. These extensive benchmarking analyses demonstrated that SpaHDmap generates highly interpretable, high-resolution embeddings and helps uncover complex and fine spatial structures that were challenging for available methods.

## Results

### SpaHDmap fuses expression and histology image for high-resolution dimension reduction

SpaHDmap utilizes a multi-modal neural network that takes advantage of the high-dimensionality of transcriptomics data and the high resolution of histology image data to achieve interpretable high-resolution dimension reduction (Fig.1; Methods). The high-dimensional gene expression data enable refined functional annotations, while the histology image data help to enhance spatial resolution. After embedding learning, SpaHDmap can perform high-resolution downstream analyses, such as embedding-associated gene identification, high-resolution spatial domain detection, and gene expression recovery.

**Fig. 1.**
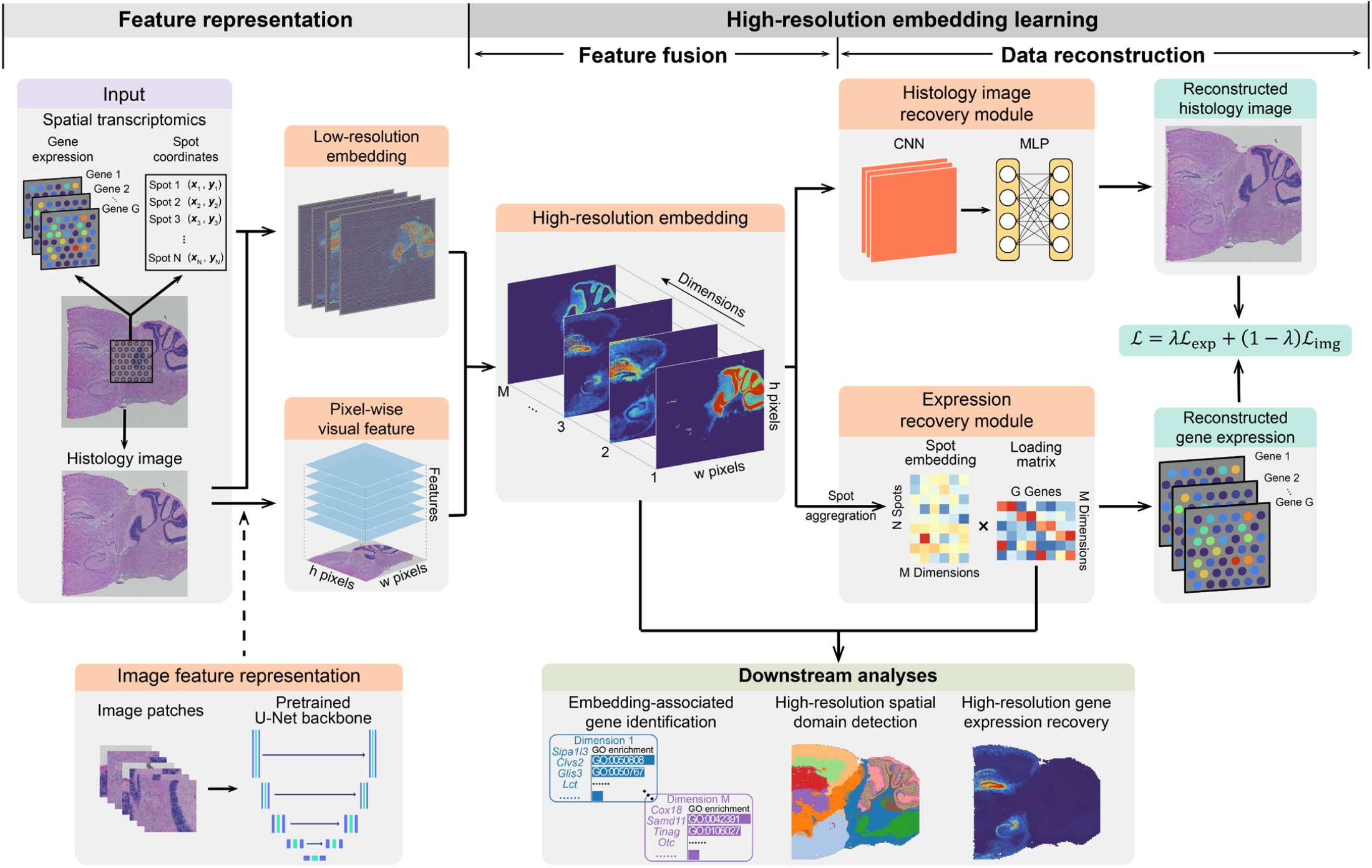
Overview of SpaHDmap. SpaHDmap consists of two components: the feature representation and high-resolution embedding learning components. In the feature representation component, SpaHDmap independently learns a low-resolution embedding from the ST data and a pixel-wise feature representation from the histology image data. In the high-resolution embedding learning component, SpaHDmap integrates the low-resolution embedding and the pixel-wise image features to learn a high-resolution embedding. This is achieved by reconstructing both the gene expression data and the image data from the learned embedding using an encoder-decoder framework. With the learned embedding, SpaHDmap can perform various downstream analyses, including identification of genes associated with the learned embeddings, high-resolution spatial domain detection, and gene expression recovery. CNN: convolutional neural network. MLP: multilayer perceptron.

The SpaHDmap framework comprises two major components: feature representation and high-resolution embedding learning (Fig.1). (1) In the feature representation component, SpaHDmap independently learns a low-resolution embedding from the ST data and a pixel-wise visual feature from the image data (Fig.S1; Methods). The low-resolution embedding is learned by NMF coupled with a denoising graph convolutional network (GCN). For the image feature representation, SpaHDmap pretrains a multi-channel U-Net that recovers the image patches and then transfers its backbone to extract pixel-wise visual features. (2) SpaHDmap then leverages a multi-modal fusion encoder-decoder to learn a high-resolution embedding via simultaneously reconstructing the ST gene expression and histology images. The encoder (feature fusion module) fuses and maps the low-resolution embedding and the pixel-wise image features into a high-resolution embedding space. Then, the decoder (data reconstruction module) reconstructs gene expression and histology images based on the learned embedding. Similar to NMF, the entries of the embedding and the learnable loading matrix in the expression recovery module are all non-negative, thus allowing an interpretable parts-based high-resolution representation of the data. The reconstruction loss is a weighted sum of the losses for the gene expression and histology image reconstruction. Details of the SpaHDmap framework are provided in Methods.

### SpaHDmap gives more accurate embeddings and spatial clustering in simulation

We evaluated the performance of SpaHDmap in comparison with available NMF-based methods, including NMF^29^, NSFH^15^, and SpiceMix^16^, as well as Giotto^17^, an interpretable dimension reduction method based on gene co-expression modules, using two simulation studies. Since these methods can only produce low-resolution embeddings, we thus included a high-resolution embedding method, TESLA+NMF, which employed TESLA^25^ to recover high-resolution expression from images and then applied NMF to the imputed expression data to obtain high-resolution embeddings.

The first simulation utilized an H&E image of a mouse cerebellum for data generation (Fig.2a; Methods). The image was segmented into three regions based on manual annotation, and intensities for the embedding dimension of each region were generated. Gene expression at individual generated spots was simulated using the zero-inflated Poisson distributions, with parameters dependent on the spot’s embedding intensities. We simulated different resolutions of ST data by randomly excluding spots or adjusting the spot radius. Additionally, we introduced additive noise or dropouts to simulate different levels of noise in the ST data (Methods).

In terms of the recovery of embedding intensities, SpaHDmap consistently demonstrated superior performance, often by a large margin compared to other methods, as measured by the mean absolute error (MAE) between the true and inferred embedding intensities (Fig. 2b; Fig. S2a, c-e; Methods). After obtaining the embedding, we proceeded to identify spatial domains by applying the K-means algorithm to the learned embedding and compared clustering accuracy using the Adjusted Rand Index (ARI) (Methods). Again, SpaHDmap achieved substantially higher ARIs than other methods in all simulation scenarios (Fig. 2c; Fig. S2b), demonstrating the effectiveness of SpaHDmap’s integration of ST expression data and histology images. Figures 2d and 2e show the local embedding and clustering result from a simulation dataset (spot radius: 107 pixels) within a refined region of interest (ROI) (Fig. 2a). Even though the resolution of ST expression data was low, the spatial fine structures could still be faithfully captured by SpaHDmap’s embedding (Fig. 2d), leading to significantly more accurate clustering than other methods (Fig. 2e), especially in the sub-regions with very fine structures (light blue boxes in Fig.2d,e).

**Fig. 2.**
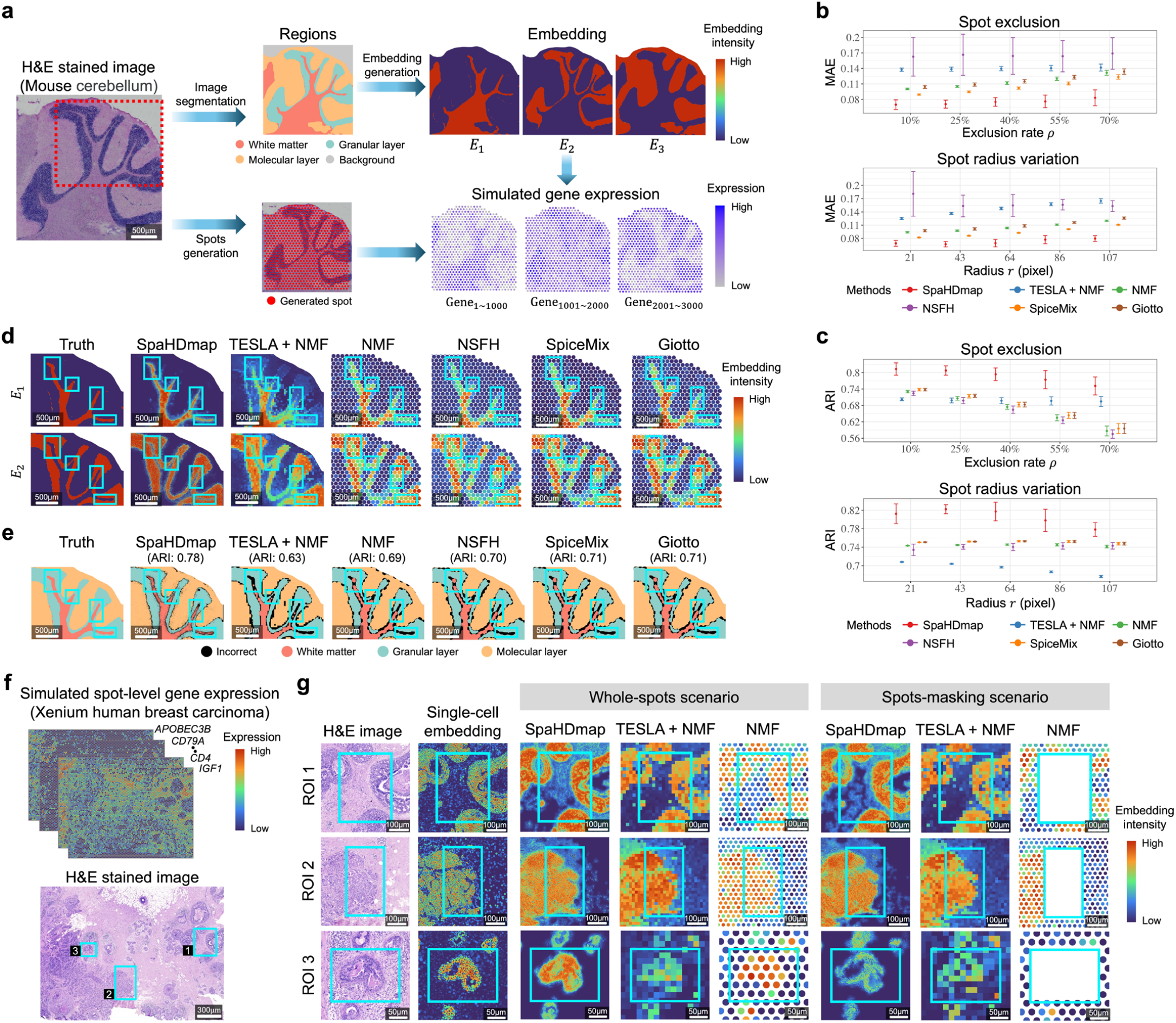
Comparison of SpaHDmap with interpretable embedding learning methods using simulation. **a**. Generation of the first simulation dataset. The H&E image of a mouse cerebellum sample is manually segmented into three brain regions and a background region. Spot locations are randomly generated on the non-background area of tissue. Each brain region is then associated with a specific embedding dimension, and spot-wise ST data are generated based on the location of sequenced spots and the embedding dimensions. Scale bar: 500 μm. **b-c**. Error bar plots showing (b) the MAEs between the true and inferred embeddings given by different methods under various simulation scenarios, and (c) the ARIs between clustering results given by different methods and the true spatial domains. Error bars represent the mean ± standard deviation (SD). **d-e**. Zoomed-in views of the red-box region highlighted in panel (a). Scale bar: 500 μm **d**. The local true embeddings E1 and E2 and corresponding embeddings estimated by different methods. Spatial intensities of each embedding dimension are linearly scaled to [0,1] for visualization. **e**. The local clustering results given by different methods. The numbers on the top of the subfigures are the ARIs in this local region. Black color represents regions that are incorrectly clustered. **f**. Heatmaps showing examples of the second simulation data generated from a human breast carcinoma sample and the corresponding H&E stained image. Scale bar: 300 μm. **g**. Zoomed-in views of the inferred embedding dimensions learned by various methods from one of the second simulation datasets (spot radius: 30 pixels) in the three selected ROIs in panel (f). Scale bar of ROI 1-3: 100, 100, and 50 μm. The whole-spots scenario means that expression of all spots of the simulation dataset are used for embedding learning, and the spots-masking scenario means that the ST expressions in the light blue boxes are masked for embedding learning.

The second simulation generated synthetic data from a human breast cancer dataset sequenced by 10X Xenium^3^, which comprises single-cell expression data for 313 genes. We aggregated expressions from neighboring single-cells to create low-resolution spot-level ST data with varying spatial resolutions (Fig. 2f; Fig. S3a, b; Methods). Comparing the embeddings given by different methods with the single-cell-level NMF embedding from the original Xenium dataset (Fig. S3a), SpaHDmap consistently produced more accurate embeddings (Fig. S3c). To further assess SpaHDmap’s capability in recovering fine structures, we conducted a challenging simulation scenario by excluding spots in three ROIs that contained local fine structures (Fig. 2g; Fig. S3d). Although several methods could recover these local patterns when the spots in the ROIs were not excluded, only SpaHDmap and TESLA+NMF were able to provide embeddings for all three ROIs in this challenging scenario (Fig. 2g; Fig. S3e). Notably, only SpaHDmap could recover the local pattern for the extremely refined ROI 3.

### SpaHDmap robustly recovered interpretable high-resolution embeddings in mouse brain datasets

We applied SpaHDmap to the 10X Visium ST dataset MBC-01 from an adult mouse brain coronal section, comprising three immunohistochemistry (IHC) stained images (Fig. S4a; Methods). The embeddings identified by SpaHDmap were enriched in known specific brain regions (Fig. S4). For example, E15 was mainly enriched in isocortex layers 1, 2/3 and 4, E18 in isocortex layers 4 and 5, and E9 in isocortex layers 6a and 6b, E10 in the dentate gyrus (DG), E14 in the pyramidal layers CA1 and CA2, E16 in the pyramidal layers CA2 and CA3 (Fig. 3a, b). Many of these embeddings were also identified by other methods (Fig. S5). However, most embeddings given by other methods were of low resolution. TESLA+NMF could give high-resolution embeddings, but several of its embeddings did not correspond well to the known brain regions (e.g., TESLA+NMF’s embedding dimensions E1 and E18 in Fig. S5). Zoomed-in views of SpaHDmap’s embedding dimensions E1 and E6 in four selected ROIs showed that local fine structures unveiled by these embeddings were well-supported by staining for neuron-specific nuclear binding protein (NeuN) or glial fibrillary acidic protein (GFAP) (Fig. 3c,d). In comparison, TESLA+NMF could only identify the local fine structure in ROI 1, and other methods could hardly detect any due to their low resolution (Fig. S6).

**Fig. 3.**
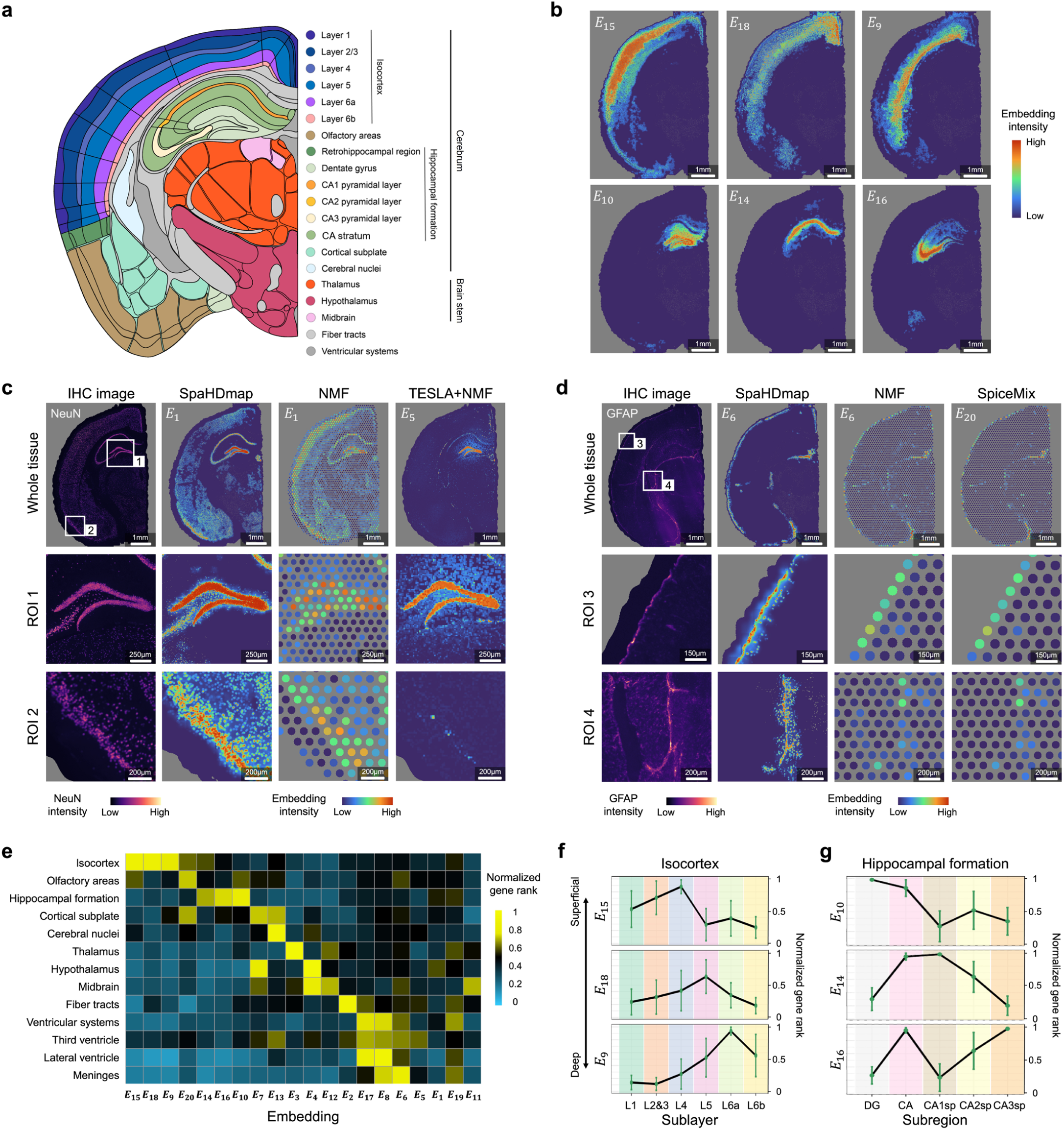
Embeddings learned from the adult mouse brain coronal section data MBC-01. **a**. Anatomical annotations from the Allen Reference Atlas – Mouse Brain. **b**. Representative embedding dimensions identified by SpaHDmap including the isocortex-associated dimensions (E15, E18, E9) and hippocampal-formation-associated dimensions (E10, E14, E16). **c-d**. Embedding dimensions associated with NeuN or GFAP protein expressions inferred by SpaHDmap, TESLA+NMF, NMF and SpiceMix and their zoomed-in views in four selected ROIs. Scale bar of whole image: 1mm. Scale bar of ROI 1-4: 250, 200, 150 and 200 μm. **e**. A heatmap showing the mean rankings of brain-region marker genes in each embedding dimension. Gene rankings are normalized to [0,1]. In each embedding dimension, the genes are ranked increasingly by their corresponding values in the loading matrix. **f-g**. Error bar plots showing gene rankings of marker genes of subregions of isocortex and hippocampal formation in associated embedding dimensions. Error bars: median ± median absolute deviation (MAD, a robust SD). DG: dentate gyrus; CA: cornu ammonias; CA1sp: pyramidal layer of field CA1; CA2sp: pyramidal layer of field CA2; CA3sp: pyramidal layer of field CA3.

To further investigate the enrichment of embedding dimensions in specific brain regions, we obtained genes upregulated in different brain regions from the Allen Mouse Brain Atlas^30^ and investigated their rankings in SpaHDmap’s embedding dimensions (Methods; Supplementary Table 2). The marker genes of different brain regions ranked high in only a few embeddings, implying that the embeddings were associated with specific brain regions (Fig. 3e). For example, marker genes of olfactory areas were only enriched in embedding E20, those of cerebral nuclei in E13, and those of thalamus in E3. Isocortex markers ranked high in three embedding dimensions (E15, E18 and E9). We further obtained marker genes of the sublayers of the isocortex and found that isocortex layers 1, 2&3 and 4 were enriched in embedding E15, layer 5 in E18 and layer 6a/6b in E9 (Fig. 3f), demonstrating that E15, E18 and E9 were associated with the sublayers of the isocortex. Similarly, hippocampal formation markers were enriched in three embeddings (E10, E14 and E16), and these embeddings were associated with subregions of the hippocampal formation (Fig. 3g).

Next, we applied SpaHDmap to another two mouse brain coronal sections, MBC-02 and MBC-03, where MBC-02 contained two IHC images and MBC-03 contained one H&E image (Methods; Fig. S7a). Note that the image data of these two datasets differed from the image data of the previous MBC-01 dataset (Fig. S7a). We found that the embeddings learned from the MBC-02 and MBC-03 datasets aligned well with the embeddings learned from the MBC-01 dataset at high resolution (Fig. S7b), and their gene ranks were also highly correlated with embeddings learned from the MBC-01 dataset (Fig.S8a, b; Supplementary Table 2). Gene enrichment analysis further confirmed that the embeddings learned from the MBC-02 and MBC-03 datasets were associated with specific brain regions (Fig.S8c-f). In addition, we analyzed two adjacent mouse posterior brain sagittal section datasets (MPBS-01 and MPBS-02) sequenced by the 10X Visium with H&E images (Methods; Fig.S9a). Again, both slices consistently exhibited enrichment of embeddings in specific brain regions (Fig.S9b, c), marker genes of the brain regions consistently ranked high in their respective embeddings (Fig.S9d; Supplementary Table 2), and the embeddings learned from both slices showed strong correspondence (Fig.S9e). These results demonstrate that SpaHDmap can robustly identify high-resolution, interpretable embeddings from ST data with various types of histology images.

### SpaHDmap identified fine-grained spatial structures in mouse brain data

Using the embeddings from the mouse brain dataset MBC-01, we performed clustering analysis to detect spatial domains (Fig.4; Fig.S10; Methods). For comparison, we further included three state-of-the-art spatial domain detection methods, iStar, SpaGCN and stLearn, that also utilized images. SpaHDmap, TESLA+NMF, and iStar achieved the highest spatial resolution in spatial domain detection. Although SpaGCN and stLearn also leveraged image data, their spatial resolutions still remained at the spot level. Overall, spatial domains identified by SpaHDmap showed better concordance with known mouse brain domains^31^ compared to other methods, and this concordance was robust to the resolution parameter of the Louvain clustering algorithm (Fig. 3a; Fig.S10).

The advantage of SpaHDmap was particularly evident in the hippocampal formation region that contained fine-grained spatial structures (Fig. 4a, b). Only SpaHDmap successfully identified and clearly delineated subregions of hippocampal formation and their complex topological relationships (Fig. 4a, b). Among the low-resolution methods, stLearn and SpiceMix performed the best in detecting these local subregions. However, due to their lower resolution, the complex topological relationships between the subregions identified by these methods were much less clear. The high-resolution methods, iStar and TESLA+NMF, largely recovered the pyramidal layers CA1, CA2 & CA3, and the granule cell layer of the dentate gyrus (DG). However, iStar incorrectly separated pyramidal layer of CA1 and the granule cell layer of DG into two clusters each, TESLA+NMF misclassified many points in pyramidal layers to another cluster, and both methods merged stratum layers CA1, CA2 & CA3 and the molecular layer DG into a single cluster (Fig. 4a). Known marker genes^32, 33^ of the hippocampal pyramidal layers CA1, pyramidal layers CA2 and CA3, and the granule cell layer DG were highly expressed in these three subregions identified by SpaHDmap (Fig. 4c; Fig. S11a). Moreover, spatial expression patterns of these genes were clearer in the recovered high-resolution expression by SpaHDmap than in the raw expression (Fig. 4c; Fig. S11).

**Fig. 4.**
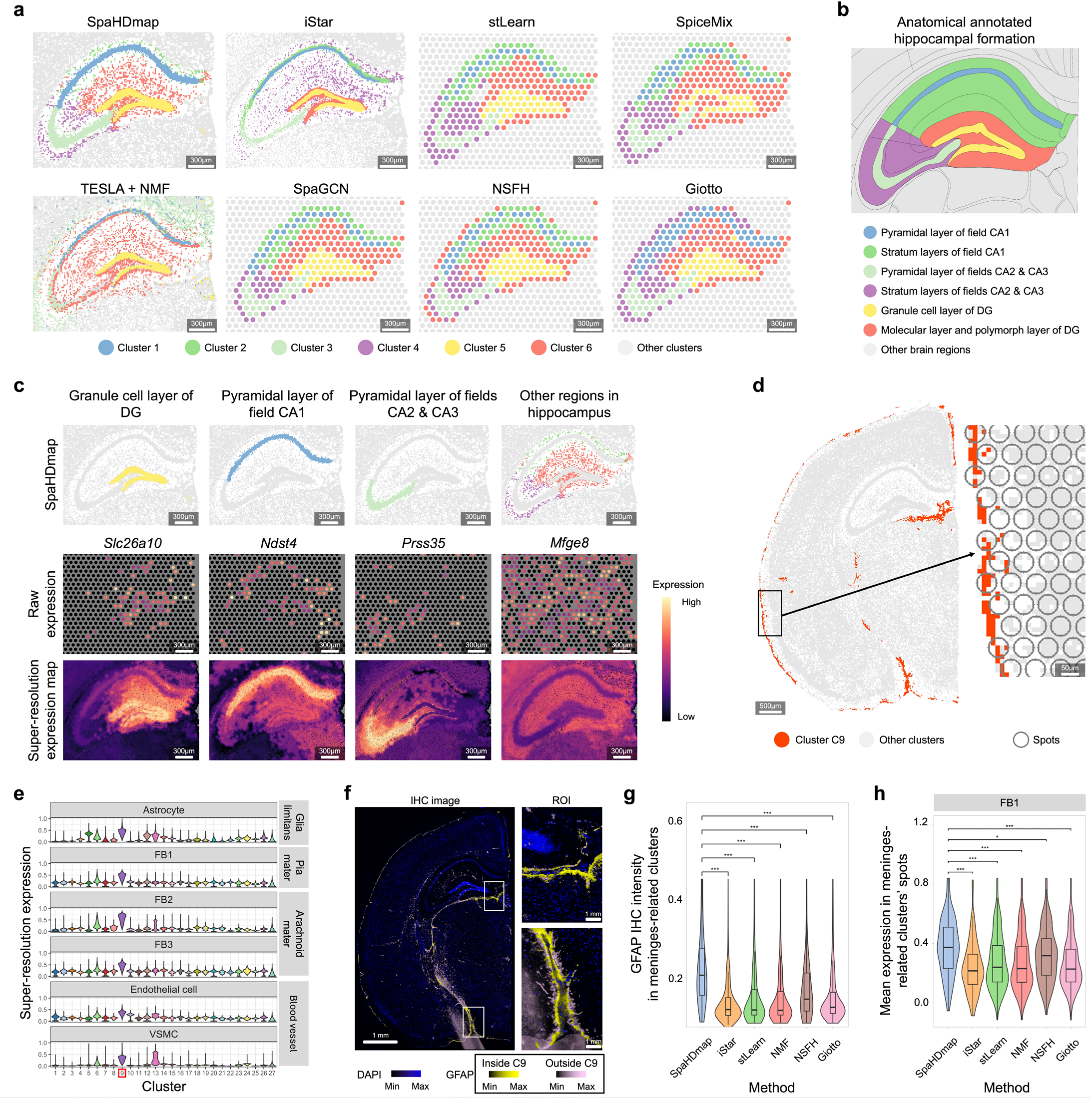
Spatial domains detected from the adult mouse brain coronal section data MBC-01. **a**. Zoomed-in views of clusters identified by different methods in the hippocampal formation. The spatial clusters were generated by Louvain clustering with resolution 2.0 on the embeddings learned by different methods, except for iStar which used the same number of clusters as SpaHDmap. For spot-level methods, spot radii are enlarged 1.2-fold for visualization. Scale bar: 300 μm. **b**. Anatomical annotations from the Allen Reference Atlas – Mouse Brain hippocampal formation. DG: dentate gyrus; CA: cornu ammonias. **c**. The raw expression and high-resolution expression recovered by SpaHDmap for the marker genes of hippocampal formation subregions. Scale bar: 300 μm. **d**. A refined cluster C9 located around the meninges. Scale bar: 500 μm. Right panel: The zoomed-in view of an ROI showing that this cluster is often smaller than a spot. Scale bar: 50 μm. **e**. Violin plots showing the recovered high-resolution expression of marker genes of major cell types in glial limitans and leptomeninges across different spatial clusters. FB: meningeal fibroblasts subtype. VSMC: vascular smooth muscle cell. **f**. The IHC image of GFAP and 4’,6-diamidino-2-phenylindole (DAPI). Blue: DAPI; Yellow: GFAP in cluster C9; Pink: GFAP in other clusters. Right panels: Zoomed-in views of two ROIs. Scale bar: 1 mm. **g**. Violin plots of the IHC intensity levels of GFAP in meninges-related clusters identified by different methods. **h**. Violin plots of mean expressions of FB1 marker genes in meninges-related clusters’ spots identified by different methods. P-values are obtained using the two-sided Wilcoxon rank-sum test. *: p-value ≤ 0.05; ***: p-value ≤ 0.001.

SpaHDmap detected a thin cluster C9 located at the meninges (Fig. 4d; Fig. S12a). The meninges are important protective membranes that envelop the brain and consist of three layers: dura, arachnoid, and pia mater^34, 35^. The arachnoid and pia mater together form the leptomeninges. Beneath the pia mater lies the glia limitans, a thin membrane layer primarily composed of astrocyte end-feet^36^. Within the meninges, a network of cerebral arteries and veins traverses inward to the brain^37, 38^. Cluster C9 highly expressed marker genes of astrocytes and major cell types in the leptomeninges^36^, including meningeal fibroblasts subtype 1 (FB1) in the pia mater, FB2, FB3 in the arachnoid mater, as well as endothelial cells and vascular smooth muscle cells (VSMCs) within blood vessels, but not marker genes of cell types in the dura mater (Fig. 4e; Fig. S12b,c). GFAP, the protein marker of astrocytes, showed a higher IHC intensity in cluster C9 than in other clusters (Fig. 4f). These data suggested that cluster C9 was composed of the leptomeninges and the glia limitans. Several other methods also detected clusters located near the meninges (Fig. S12d). However, cluster C9 was much thinner and exhibited higher IHC intensity of GFAP (Fig.4g) as well as higher expression levels of FB1, FB2, and endothelial cell markers than other methods (Fig. 4h; Fig. S12e), demonstrating that C9 matched better to the thin structure of the leptomeninges and glia limitans. These results highlight SpaHDmap’s capability to enhance embedding resolution and capture fine-scale spatial structures.

### SpaHDmap recovered condition-specific embeddings across multiple tumor samples

We applied SpaHDmap to a 10X Visium dataset of sonic hedgehog (SHH) medulloblastoma derived from PDOX mouse models^39^. The dataset consisted of two Palbociclib-treated samples (Palbociclib A and B) and two untreated samples (Control C and D) (Fig. 5a). Joint analysis of these four samples using SpaHDmap yielded 20 embedding dimensions (Methods): ten enriched in normal mouse brains, five in implanted tumors and five in tumor-normal interfaces, as indicated by their respective high intensity at the mouse, human and human-mouse mix spots, as well as their rankings of mouse and human genes (Fig. 5a, b; Fig. S13; Supplementary Table 2). The normal embeddings aligned well with known mouse brain structures, such as the Purkinje layer (E4), granular layer (E8), and molecular layer (E9) in the cerebellar cortex (CBX) (Fig. 5b; Fig. S13). Astrocytes and microglia (resident macrophages in brains) have been reported to be enriched at the tumor-mouse brain interface^39^. Consistently, the interface embedding dimensions E11 and E15 were enriched with marker genes of astrocytes and microglia, respectively (Fig. 5b; Fig. S13).

**Fig. 5.**
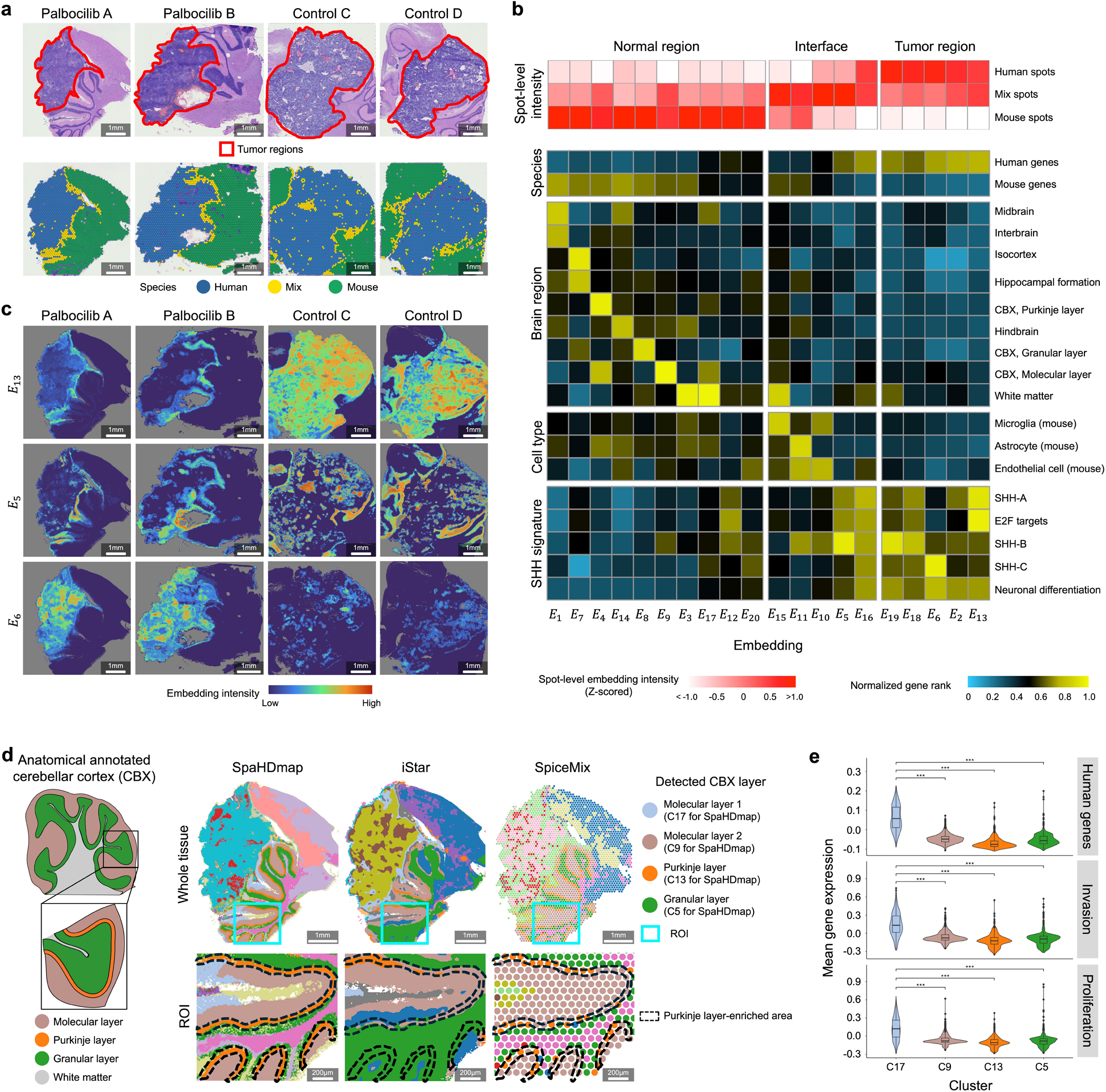
Embedding and clustering results of the four PDOX samples. **a**. Upper panel: The H&E images with annotated tumor regions. Lower panel: Spot locations along with their species classifications based on their expressions of human and mouse genes. Scale bar: 1mm. **b**. Rows 1-3 showing the 95th percentile of spot-level embedding intensity for different species and subsequent rows displaying the mean rankings of various gene signatures across each embedding dimension. Gene rankings were normalized to [0,1]. CBX: cerebellar cortex. **c**. The embedding dimensions (E6, E13, E5) associated with SHH transcriptional signatures. Scale bar: 1mm. **d**. Left panel: Anatomical annotations from the Allen Reference Atlas – Mouse Cerebellum Cortex. Right panel: Spatial clusters identified by SpaHDmap, iStar, and SpiceMix in the Palbociclib A section. For SpiceMix, its filtered low-quality spots are colored in grey, and spot radius is enlarged 1.3-fold for better visualization. Scale bar: 1mm. Bottom panels: Zoomed-in views of the ROI in the light blue box of the top panel. Scale bar: 200 μm. **e**. Violin plots of the mean normalized expressions of human genes, tumor invasion genes, and tumor proliferation genes across spots in clusters identified by SpaHDmap. P-values are obtained using the two-sided Wilcoxon rank-sum test. ***: p-value ≤ 0.001.

Hovestadt et al.^40^ identified three important transcriptional signatures in SSH medulloblastoma: cell cycle activity (SHH-A), undifferentiated progenitors (SHH-B) and differentiated neuronal-like programs (SHH-C). We found that embedding dimension E13 was enriched with SSH-A and E2F target genes, E5 with SSH-B genes, and E6 with SSH-C and neuronal differentiation genes (Fig. 5b; Fig. S14). Notably, E6 showed increased intensities in Palbociclib-treated samples compared with untreated samples. In contrast, E13 and E5 exhibited reduced intensities in Palbociclib-treated samples, and the reduction largely only occurred in the interior regions of the tumors but not at the tumor boundaries (Fig. 5c). These results were consistent with the observation that Palbociclib could enhance neuronal differentiation and reduce cell proliferation, but have less impact at tumor boundaries^39^, and consistent with the fact that the CDK4/6 inhibitor Palbociclib can downregulate E2F-mediated transcription of cell cycle genes, thereby inducing G1 arrest and inhibiting cancer cell proliferation^41^.

Using SpaHDmap’s embeddings, we performed a joint cluster analysis of the four samples (Fig. S15a). As expected, normal brain regions shared highly similar clusters across all samples, while tumor regions were more heterogeneous. In tumor regions, samples with the same treatment had more shared clusters than samples with different treatments. Among the spatial clustering methods, SpaHDmap and SpiceMix allowed joint analysis of multiple samples, resulting in naturally aligned clusters between samples. For other methods, aligning clusters across samples was more challenging (Fig. S15a). Furthermore, SpaHDmap clearly characterized the ordered layers of the CBX (Fig. 5d; Fig. S15a-d), including the molecular layer (C17, C9), Purkinje layer (C13) and granular layer (C5). In comparison, other methods could not clearly delineate these layers, especially the fine Purkinje layer (black dotted area in Fig. 5d and Fig. S15b). Among the low-resolution methods, SpiceMix performed the best in detecting the CBX layers, possibly due to its ability to integrate multiple samples. Interestingly, the molecular layer cluster C17 detected by SpaHDmap was adjacent to tumor regions and thus more susceptible to tumor cell infiltration (Fig. S15e). Consistently, C17 was significantly enriched with human genes and exhibited higher invasion and proliferation activity compared to other normal clusters (Fig. 5e). Additionally, the C17 areas decreased in Palbociclib-treated samples compared with untreated samples, likely due to Palbociclib’s effect on reducing tumor cell proliferation and invasion (Fig. S15e). These analyses demonstrate that joint high-resolution embedding learning by SpaHDmap can effectively capture multi-sample fine-scale structures and enable comparative analysis between multiple samples.

### SpaHDmap depicts tumor heterogeneity and immune activities in CRC samples

We applied SpaHDmap to three newly profiled ST CRC samples (CRC-01, 02, 03) for spatial cluster detection. Pathologist annotations indicated that sample CRC-01 comprised four major spatial clusters: tumor, stroma, necrosis, and muscle (Fig. 6a). SpaHDmap identified 16 refined spatial subclusters within these broader categories, including 6 tumor, 7 stromal, 1 necrotic, and 2 muscle subclusters (Fig. 6b; Fig. S16a). Compared with spatial clusters given by available methods, clusters given by SpaHDmap agreed better with the H&E image (Fig. 6a-c; Fig. S16b-d; Supplementary Note 1). For example, only SpaHDmap accurately distinguished necrosis from tumor in ROI 1, and tumor from stroma in ROI 2 and ROI 3 (Fig. 6d; Fig. S16c).

**Fig. 6.**
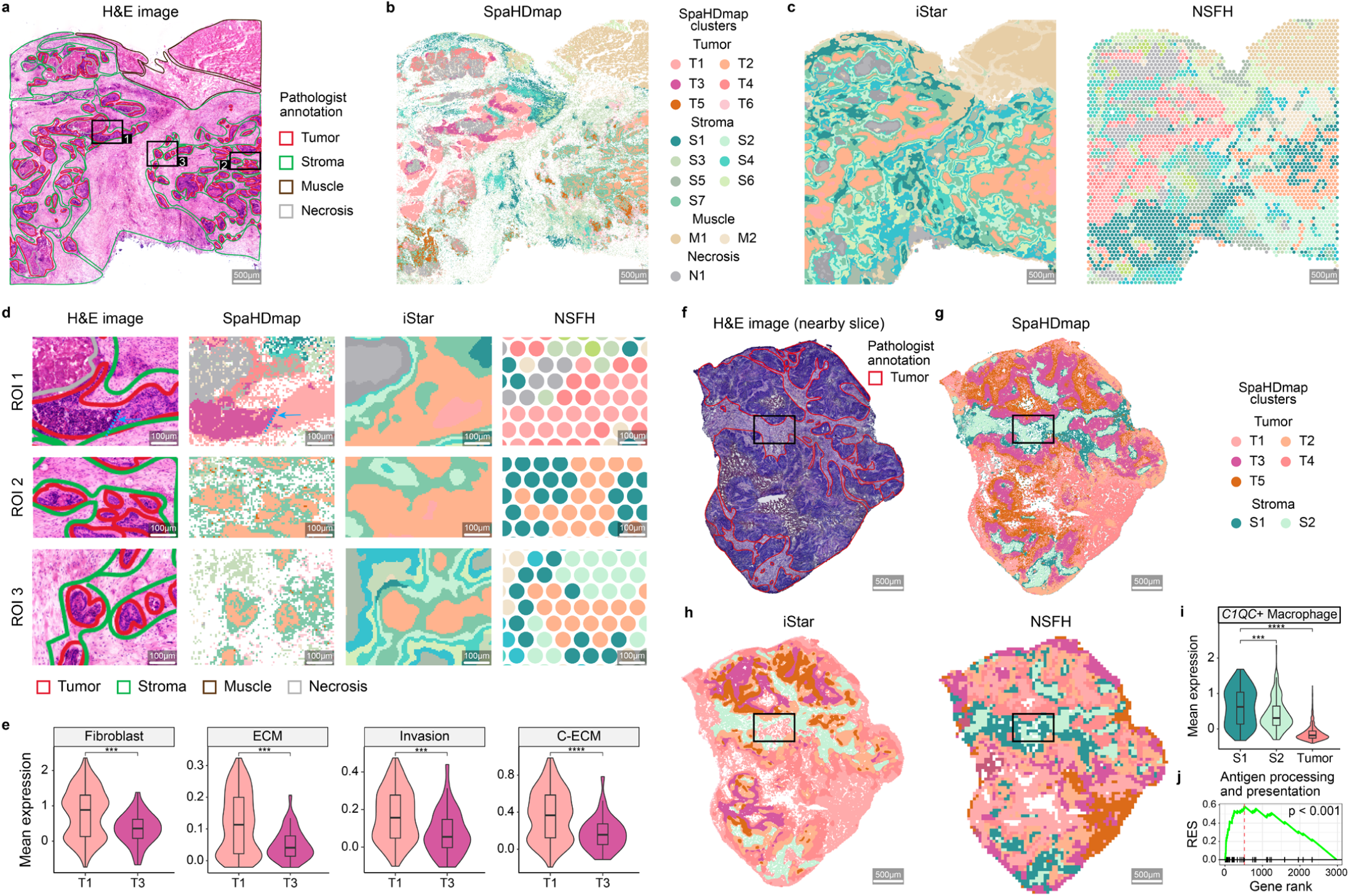
Analysis of colorectal cancer samples. **a-e**. Analysis of sample CRC-01. **a**. The H&E image of sample CRC-01 with annotation by our pathologist. Annotated tumor, stroma, muscle and necrosis regions are outlined in red, green, brown and grey, respectively. Scale bar: 500 μm. **b**. Spatial clusters identified by SpaHDmap. Spatial clusters are annotated by their marker genes (see Fig. S16a). Scale bar: 500 μm. **C**. Spatial clusters identified by iStar and NSFH. For NSFH, its filtered low-quality spots are colored in grey, and spot radius is enlarged 1.2-fold for better visualization. Scale bar: 500 μm. **d**. Zoomed-in views of H&E images and clusters identified by SpaHDmap, iStar and NSFH. ROIs correspond to black-box regions in (a). Scale bar: 100 μm. **e**. Violin plots of mean gene expression of fibroblast, extracellular matrix (ECM), cancer-invasion and cancer-associated ECM (C-ECM) signatures. **f-j**. Analysis of sample CRC-03. **f**. The H&E image of a nearby slice of sample CRC-03 with pathologist annotation. Annotated tumor regions are outlined in red. Other regions are all stroma regions. Scale bar: 500 μm. **g**. Spatial clusters identified by SpaHDmap. Scale bar: 500 μm. **i**. Spatial clusters identified by iStar and NSFH. For NSFH, low-quality spots are colored in grey, and spot radius is enlarged 1.2-fold for better visualization. Scale bar: 500 μm. **i**. Violin plots of mean gene expression of *C1QC*+ macrophage markers. P-values are obtained using the two-sided Wilcoxon rank-sum test. **j**. Gene set enrichment analysis (GSEA) on cluster S1-related genes for antigen processing and presentation. The enrichment plots contain profiles of the running enrichment scores (RES) and positions of gene set members on the rank ordered list in GSEA. ***: p-value ≤ 0.001; ****: p-value ≤ 0.0001.

Among the tumor clusters, T1, T2 and T3 resembled the CMS2 subtype of colorectal cancer, T6 was similar to the CMS4 subtype, and T4 and T5 appeared to be intermediate states between CMS2 and CMS4^42, 43^ (Fig. S17a; Supplementary Note 1), suggesting that they were distinct tumor spatial subclusters. Notably, in ROI 1, the tumor clusters T1 and T3 had a clear boundary on the H&E image (indicated by the blue line in Fig.6d), which were only detected by SpaHDmap (Fig. 6d; Fig.S16c). The H&E image showed that T1 was more enriched with fibroblast than T3 (Fig. S17b). Consistently, T1 showed significantly higher expressions of fibroblast and extracellular matrix (ECM) genes than T3 (Fig. 6e). Deconvolution analysis by RCTD^44^ (Methods) also revealed lower tumor purity in T1 (0.56) compared with T3 (0.70). Further, T1 had significantly higher cancer-associated ECM^45^ (C-ECM) and invasion scores (Fig. 6e), indicating that T1 might be more aggressive and invasive. In fact, genes upregulated in T1 relative to T3 were predictive of overall worse survival of colorectal patients in the TCGA data^46^ (Fig. S17c). Taken together, these results supported that T1 and T3 were distinct tumor clusters with important clinical implications. Similarly, SpaHDmap accurately detected fine structures in sample CRC-02 (Fig. S18; Supplementary Note 1).

Sample CRC-03 was sequenced by Stereo-seq technology without a paired histology image. To apply SpaHDmap and other image-dependent methods, we generated a pseudo-image from the noisy subcellular Stereo-seq data (Fig. S19a; Methods). SpaHDmap identified 5 tumor subclusters and 2 stroma subclusters through integrating low-resolution bin-level expression data with the pseudo-image (Fig.6f, g; Fig. S19a, b; Supplementary Note 1). Compared with other high-resolution methods iStar and TESLA+NMF (Fig. 6h; Fig. S19c), clusters given by SpaHDmap agreed better with the H&E image of a nearby tissue slice (Fig. 6f). For example, in an ROI (black-box region in Fig. 6f-h; Fig. S19c, d), SpaHDmap and spot-level methods correctly distinguished stroma and tumor regions. In contrast, iStar and TESLA+NMF misclassified large tumor regions as stroma, possibly because they were designed for integrating H&E images and may not be suitable for analyzing pseudo-image generated from Stereo-seq data.

Moreover, SpaHDmap identified a refined cluster S1 around the tumor boundary (Fig. 6g). Compared with the nearby stroma cluster S2, cluster S1 was more enriched with *C1QC*+ macrophages^47^ (Fig. 6i; Fig. S19e). *C1QC*+ macrophages were known to have increased antigen-presenting activities^47^. Consistently, genes highly related to S1 were significantly enriched with antigen-presenting genes (Fig. 6j), suggesting S1’s potential role in tumor-associated inflammation. Additionally, genes highly expressed in S1 were associated with overall better survival of colorectal patients in the TCGA dataset (Fig. S19f), indicating the potential clinical relevance of this refined cluster. These analyses demonstrate the superior performance of SpaHDmap in accurately identifying fine spatial structures of tumor samples at high resolution, even in the absence of a paired histology image.

## Discussion

In this paper, we developed SpaHDmap that generates high-resolution, interpretable embeddings and identifies spatial structures by effectively integrating ST gene expression data with high-resolution histology images. SpaHDmap enables joint analysis of multiple samples and supports various types of histology images. This novel approach opens up new possibilities for uncovering fine-grained spatial patterns and biological insights that were previously unattainable due to the resolution limitations of ST technologies. Extensive benchmarking analyses using simulated and real ST datasets demonstrated that SpaHDmap consistently outperformed state-of-the-art methods in embedding learning and spatial domain detection, particularly in challenging scenarios with complex spatial structures. SpaHDmap is comparable to available methods in terms of computational speed, but requires slightly more memory usage (Fig. S20).

Deep learning methods have been widely used for dimension reduction or clustering analysis of single-cell RNA-seq and ST data, often outperforming traditional statistics and machine learning methods^48-51^. However, their black-box nature often hinders the interpretability of the learned features, which is crucial for understanding important structures in high-dimensional single-cell RNA-seq or ST data. To address this limitation, SpaHDmap introduces a framework that fuses NMF with a deep neural network, enabling interpretable dimension reduction while leveraging the power of deep learning and rendering higher resolution. As demonstrated in applications to mouse brain and PDOX datasets, the learned embeddings, along with the associated genes, represented unique spatial expression patterns or known tissue structures, and could even reveal important transcriptional signatures related to tumor treatment, thus facilitating a more straightforward data interpretation. The NMF component can also be replaced with other spatially-aware dimension reduction methods, such as NSFH^15^ or SpatialPCA^13^, which might enable more accurate embedding estimation with spatial patterns.

Based on the high-resolution embeddings, SpaHDmap enables the identification of refined spatial structures. In the mouse brain data, SpaHDmap identified fine-grained structures in hippocampal formation and the thin structure of the leptomeninges and glia limitans (cluster C9). Beyond physical protection, the meninges play important roles in central nervous system homeostasis, development, immunity, and pathology^34, 37^. Accurate meningeal detection is crucial for understanding their diverse functions and involvement in brain health and disease. SpaHDmap’s ability to resolve the intricate anatomy of leptomeningeal and glia limitans highlights its potential for enhancing the study of these important delicate brain structures. One potential limitation is that SpaHDmap could not separate the two sublayers of the leptomeninges and the glia limitans, as the pia mater and the glia limitans are extremely thin, comprising only one or two layers of cells, making their delineation highly challenging.

In CRC-01, we identified two spatially adjacent tumor clusters, T1 and T3, with significantly different levels of invasiveness and fibroblast infiltration, despite having similar copy number profiles (Fig. S17e). This suggests that the difference in invasiveness between T1 and T3 is more likely associated with their varying fibroblast infiltration levels rather than their genetic differences. Cancer-associated fibroblasts (CAFs) can modify ECM proteins to produce stiff and intricated fiber organization, thus increasing invasion potential and may even hinder drug delivery^52^. The presence of adjacent tumor clusters like T1 and T3, with differences in fibroblast infiltration, highlights the potential impact of CAFs on the local tumor microenvironment and the resulting heterogeneity in invasiveness. Analyzing cohorts with larger samples using SpaHDmap could further explore the prevalence and significance of adjacent tumor clusters with distinct fibroblast infiltration levels and their association with various tumor phenotypes.

In CRC-03, we identified a stroma cluster S1 near the tumor boundary that was enriched with *C1QC*+ macrophages. Previous studies have shown that *C1QC*+ macrophages are pro-inflammatory, and interact with other immune cells, particularly playing a role in recruiting or activating T cells^47^. However, we did not observe elevated expression of T cell genes in S1, suggesting that *C1QC*+ macrophages’ role in T cell recruitment or activation might depend on additional factors in the tumor microenvironment. Consistently, *C1QC*+ macrophages could be identified in healthy individuals, again indicating that their presence alone may not be sufficient to trigger a robust T cell response^53^.

SpaHDmap could be further improved in several directions. First, SpaHDmap currently requires matched ST gene expression and histology images. However, certain ST technologies, such as Stereo-seq, do not provide paired data from the same tissue slice. Extending SpaHDmap to handle ST data with images from adjacent tissue slices would broaden its applicability. This extension would involve simultaneous embedding learning and alignment of adjacent slices. Second, with the rapid development of ST technologies, ST datasets are becoming increasingly larger. There is an urgent need for computational methods that can effectively handle large-scale ST datasets. By leveraging information from large-scale samples, such methods could enable the identification of important spatial structures or patterns that are hard to detect with only a few samples. Third, the interpretability of SpaHDmap is currently achieved by extracting gene signatures for each embedding dimension, which may represent complex biological processes. Incorporating prior knowledge of known biological pathways into SpaHDmap would enhance interpretability by breaking down these multifaceted processes into specific pathway-related gene signatures.

## Methods

### SpaHDmap algorithm

#### Data preprocessing

SpaHDmap takes spatial transcriptomics (ST) data and histology images as input. Let Ω = {1,2, … ,h} × {1,2, … , *w*} be the sequenced tissue with *h* × *w* pixels (height *h*, width *w*), *I* ∈ ℝ^*h*×*w*×*c*^ be the corresponding normalized image data (scaled to [0,1]) with *c* channels within the sequenced tissue, and 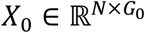 be the unique molecular identifier (UMI) count data with *G*_0_ genes at *N* sequenced spots. For spot *n* ( *n* = 1, 2, … , *N* ), let *S*_*n*_ = {(*x, y*) ∈ Ω : (*x, y*) belongs to spot *n*} stands for the set of pixels located in this spot. For each spot, the count expression data are normalized by dividing the total UMI count of the spot, multiplying 10,000, and then transforming to the natural-log-scale. After normalization, we select the top *G* spatially variable genes (default *G* = 3000) with the highest Moran’s I^54^ for downstream analyses. The resulting gene expression matrix, *X* ∈ ℝ^*N*×*G*^, and image *I* are used as the preprocessed input for SpaHDmap.

#### Image feature representation learning via U-Net-SpaHDmap

SpaHDmap pretrains a neural network, U-Net-SpaHDmap, for ST image feature representation learning based on the U-Net architecture, a widely used architecture for semantic segmentation^55^ and image restoration^56^. Specifically, U-Net-SpaHDmap first encodes 256 × 256 image patches *I* ∈ ℝ^256×256×*c*^, centered on given pixels from the ST images, using a series of convolution and down-sampling operations to extract multi-scale image features. This process results in a semantically rich representation 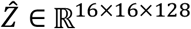. Next, U-Net-SpaHDmap uses a series of convolution and up-sampling operations to gradually merge the multi-scale image features into a pixel-wise image feature representation *Z* ∈ ℝ^256×256×32^ . Finally, a convolution operation and an element-wise Sigmoid function is used to reconstruct the image patches 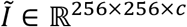 . The training data is taken as sliding overlapping 256 × 256 image patches from the ST images (15% overlap between neighboring patches). The reconstruction loss is the mean squared error (MSE) between the input and the reconstructed images, minimized by the Adam optimizer with a weight decay of 1 × 10^−5^. The initial learning rate is set as 4 × 10^−4^ and decayed to 1 × 10^−6^ using the cosine annealing strategy^57^. The mini-batch size is 32, with 5000 iterations by default.

#### Low-resolution embedding learning

SpaHDmap first learns a denoised, low-resolution embedding from the ST expression and image data. Let *M* > 0 be the dimension of the embedding space. By applying NMF to the normalized gene expression data *X* ∈ ℝ^*N*×*G*^ , we obtain an interpretable embedding of the expression data. That is, we derive a factor matrix (i.e., embedding) *F* ∈ ℝ^*N*×*M*^ and a loading matrix *W* ∈ ℝ^*M*×*G*^ (*F* ≥ 0, *W* ≥ 0), such that

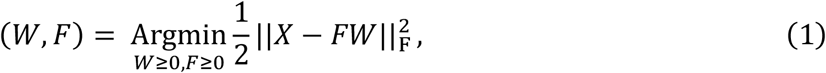

where ‖⋅‖_F_ represents the Frobenius norm. However, standard NMF can only provide embeddings for the sequenced spots, which are spatially sparse, and cannot provide any information for other locations. Moreover, this basic NMF embedding relies solely on the noisy expression data, neglecting spatial and image information, making it prone to noise and less effective at capturing fine spatial structures. To address these limitations, SpaHDmap integrates expression data, image data and spatial information using the following procedure to learn a denoised, low-resolution embedding that covers the entire sequenced tissue (Fig. S1).

In NMF, each column *F*_*j*_ (*j* = 1, ⋯ , *M*) of the factor matrix *F* represents a noisy estimate of the spatial expression pattern *j* at the *N* sequenced spots. Note that the spatial expression pattern *j* corresponds to embedding dimension *j* . We aim to denoise *F*_*j*_ and recover the low-resolution expression pattern *j* across all locations within the sequenced tissue. This is achieved using a two-step procedure: firstly, we denoise and recover the expression patterns for locations that are much denser than the sequenced spots (denoise-and-recovery step), and secondly, we extend these expression patterns to the entire sequenced tissue by Voronoi tessellations^58^ and rasterization (extension step).

The denoise-and-recovery step uses a graph convolutional network (GCN). We first construct an attributed graph *G*(*V, E, H*) , where *V* represents vertices, *E* represents edges, and *H* represents vertex attributes. The vertex set *V* consists of all *N* sequenced spots and *R* randomly sampled pseudo-spots (default: *R* = 5*N*). The attribute *H*_*v*_ ∈ ℝ^*M*^ represents expression patterns at spot *v*. For a sequenced spot *v, H*_*v*_ is the corresponding row of the factor matrix *F*. For a pseudo-spot *v*^′^, *H*_*v*′_ is unknown and needs to be learned. Edges are constructed based on spatial proximity and nearby image similarities between spots. More specifically, for each sequenced spot *v* ∈ *V*, we first identify *K*_0_ sequenced spots that are spatially closest to *v* (excluding *v* itself), and then connect *v* to the top *K* of these *K*_0_ spots whose nearby images are the most similar to the image near *v* (default: *K* = 2, *K*_0_ = 50). Similarly, the same procedure is used to connect *v* to *K* neighboring pseudo-spots. Thus, each sequenced spot is connected to *K* sequenced spots and *K* pseudo-spots. To measure the similarity between images near spots, we extract the semantically rich image representation 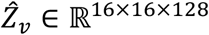 by passing the square images (256 × 256 pixels) centered at each spot through the pretrained U-Net-SpaHDmap model. Then, by averaging 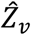 across all pixel locations 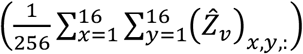, we obtain a 128-dimensional image representation for each spot and use the Pearson correlation to measure the image similarity.

Let *A* be the adjacency matrix of the graph *G* (*V, E, H*) and 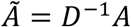 be its normalized adjacency matrix, where *D* is the out-degree matrix of *A* with 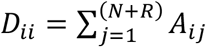. Denote 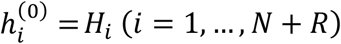 and 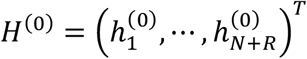. The *l*-th layer of the GCN is updated using

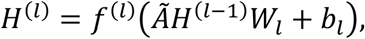

where *f*^(*l*)^ is the nonlinear activation function, *W*_*l*_ is the learnable weight matrix, and *b*_*l*_ is the learnable bias vector. The encoder of this GCN network consists of two graph convolution layers with dimension *M* → 64 → 256, while the decoder consists of two graph convolution layers with dimension 256 → 64 → *M*. The activation function for the first three layers is the rectified linear unit (ReLU), and the activation function for the final layer is the Sigmoid function. The GCN output, *H*^(4)^ ∈ ℝ^(*N*+*R*)×*M*^, is the predicted embedding for both observed spots and pseudo spots. The loss function is the MSE between predicted and input low-resolution embedding for the observed spots:

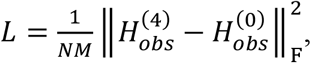

where 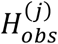 is the submatrix of *H*^(*j*)^ (*j* = 0,4) corresponding to the observed spots. An Adam optimizer, with a weight decay of 1 × 10^−5^, is used to minimize the reconstruction loss. The initial learning rate is set as 5 × 10^−3^ and decayed to 1 × 10^−3^ with the cosine annealing strategy. The number of iterations is set as 5000 by default.

Finally, we use Voronoi tessellations and rasterization to extend the learned low-resolution embedding *H*^(4)^ to all pixels in the sequenced tissue, resulting in an entire-tissue-wide low-resolution embedding *F*^low^ ∈ ℝ^*h*×*w*×*M*^ . Specifically, the entire tissue Ω is segmented into multiple multi-pixel tiles by expanding the central spatial coordinates of all observed and pseudo spots using Voronoi tessellations. Then, for the *m* -th embedding dimension ( *m* = 1,2, … , *M* ), the corresponding tissue-wide low-resolution embedding 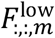 is constructed by associating the spot-wise embedding 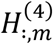 to their respective tiles.

#### High-resolution embedding learning

SpaHDmap combines the pretrained U-Net-SpaHDmap model with two modules, the feature fusion module (encoder) and the data reconstruction module (decoder), to learn high-resolution embedding. Given that ST image data are extremely high-resolution (usually with hundreds of millions of pixels), directly learning embeddings and reconstructing gene expression and image intensities for every pixel is computationally too expensive. To address this, SpaHDmap first divides the ST data into small patches, learns embeddings for each patch, and finally assembles all patches to get the entire-tissue-wide embedding. To create data patches, the sequenced tissue Ω is segmented into tissue patches Ω^(*j*)^ ⊂ Ω (*j* = 1, … , *J*) using sliding 256 × 256 rectangular with an 15% overlap between adjacent patches. Data patch 𝒟^(*j*)^ for the *j* -th tissue patch Ω^(*j*)^ (*j* = 1, … , *J*) includes the normalized expression data *X*^(*j*)^ at the spots within Ω^(*j*)^, the image data *I*^(*j*)^ ∈ ℝ^256×256×c^ and the pre-estimated low-resolution embedding *FF*^(*j*,low)^ ∈ ℝ^256×256×*M*^ within Ω^(*j*)^ , i.e., 𝒟^(*j*)^ = {Ω^(*j*)^, *X*^(*j*)^, *I*^(*j*)^,*F*^(*j*,low)^}.

For each data patch 𝒟^(*j*)^, the feature fusion module fuses the low-resolution embedding *F*^(*j*,low)^ ∈ ℝ^256×256×*M*^ with the pixel-wise image features *Z*^(*j*)^ ∈ ℝ^256×256×32^ of *I*^(*j*)^ derived from U-Net-SpaHDmap. This fusion process generates patch embedding 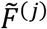, which has the same shape as *F*_(*j*,low)_:

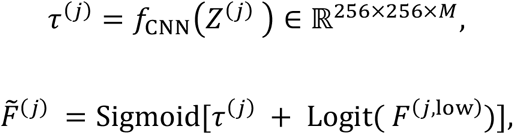

where *f*_CNN_(⋅) is a two-layer convolutional neural network (CNN), Sigmoid(⋅) and Logit(⋅) are the element-wise Sigmoid function and Logit function, respectively.

The data reconstruction module takes the high-resolution patch embeddings 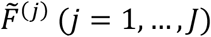 to simultaneously reconstruct both the gene expression and the image data. To reconstruct the normalized gene expression of the *n*-th sequenced spot within the *j*-th data patch 𝒟^(*j*)^ (requiring *S*_*n*_ ∩ Ω^(*j*)^ ≠ ∅), SpaHDmap first aggregates the patch embeddings 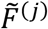 at the pixels located within the *n*-th spot to obtain spot-level embeddings, that is:

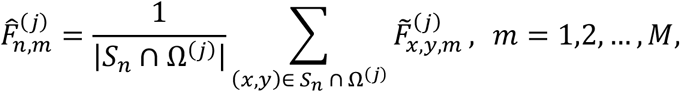

where |*S*_*n*_ ∩ Ω^(*j*)^| is the number of pixels within the intersecting area between the *n*-th spot and the *j*-th tissue patch. If the *j*-th data patch 𝒟^(*j*)^ does not contain any sequenced spot, it is simply ignored in the reconstruction of gene expression. Then, SpaHDmap reconstructs the normalized gene expression profile of the *n*-th spot in the *j*-th tissue patch using a non-negative linear combination of the spot embedding 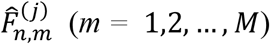 as follows:

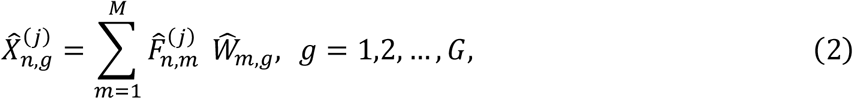

where 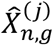 denotes the reconstructed normalized gene expression profile of the *g*-th gene at the *n*-th spot in the *j*-th tissue patch, and 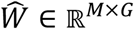 is a non-negative loading matrix that needs to be learned. The loss function for reconstructing the expression is taken as the following negative log-likelihood of Poisson distributions:

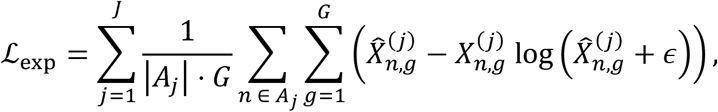

where *A*_j_ = {*n*: *S*_*n*_ ⋂ Ω ^(j)^ ≠ Ø } stands for spots belonging to the *J* -th tissue patch and ∈ = 1 × 10^-8^.

For the reconstruction of the images *I*^(*j*)^ in the *j*-th patch, SpaHDmap uses a two-layer CNN followed by an element-wise Sigmoid function to produce the reconstructed images 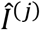 from the *j*-th patch embeddings 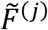 The image reconstruction loss is taken as the MSE:

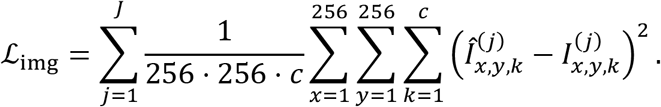

The overall reconstruction loss function is a weighted sum of the expression reconstruction loss and the image reconstruction loss

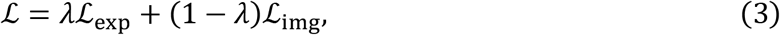

where λ ∈ [0,1] is the weight balancing the two loss functions, with a default value of 1/3. An Adam optimizer, with a weight decay of 1 x 10^-5^, is used to minimize the overall reconstruction loss. The mini-batch size is set as 32 and the number of iterations is set as 2,000 by default. During the first 1,000 iterations, parameters in U-Net-SpaHDmap are fixed, and only the feature fusion module and the image reconstruction module are updated, starting with an initial learning rate of 5 x 10^-3^ and decaying to 1 x 10^-6^ with the cosine annealing strategy. In the final 1000 iterations, all parameters in SpaHDmap are fine-tuned with a smaller initial learning rate, 4 × 10^-4^, then decaying to 1 × 10^-6^ with the cosine annealing strategy.

Finally, SpaHDmap outputs pixel-wise embeddings F^hlgh^ ∈ ℝ^*h* × *w* × *M*^ by averaging the patch embeddings 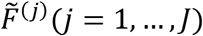 that contain th|e pixel (*x*,*y*)

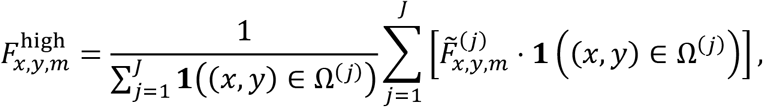

where **1** (·) is the indicator function.

### Ranking and identification of genes associated with embedding dimensions

SpaHDmap ranks genes’ contribution to each high-resolution embedding dimension 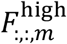 based on the *m*-th row of the non-negative loading matrix 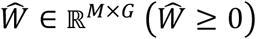 obtained from the expression reconstruction module. Specifically, the loading matrix 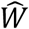 is first row-normalized to 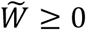with 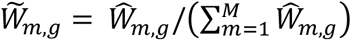. Then, for a given embedding dimension *m*, SpaHDmap ranks the genes in the decreasing order of 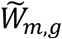 and takes the top genes as those associated with this embedding dimension.

### High-resolution spatial domain detection

SpaHDmap clusters pixels (*x, y*) based on the learned embeddings *F*^high^ ∈ ℝ^*h* × *w*× *M*^ to identify high-resolution spatial domains. Considering the extremely large number of pixels in tissue, direct pixel clustering would be computational expensive. Therefore, SpaHDmap first divides the tissue into fixed-size squares, termed as meta-pixels. Then, the pixel embeddings within each meta-pixel are averaged to produce meta-pixel embeddings, and the K-means algorithm is then applied to cluster these meta-pixels. The initial value for the K-means algorithm is taken as the Louvain clustering results based on the spot embeddings. Here, the spot embeddings are obtained using a procedure similar to that of the meta-pixel embeddings. The size of the meta-pixel is usually much smaller than the sequenced spots, and can be adjusted by users. Specifically, for the 10X Visium mouse brain coronal slices with IHC images, the size of the meta-pixel is 50 pixels, and the spot diameter is 178 pixels. For the four 10X Visium SHH medulloblastoma slices stained with H&E images, the meta-pixel size is 10 pixels, and the spot diameter is 118 pixels. For CRC-01 and CRC-02, the meta-pixel size is 20 pixels, and the spot diameter is 150 pixels. Lastly, for CRC-03, the meta-pixel size is 10 pixels, and the spot diameter is 100 pixels.

### Ranking and identification of genes associated with detected high-resolution spatial domain

Based on the *D* detected high-resolution spatial domains, we first calculate the mean embedding intensity for each spatial domain 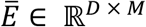, and then extract the non-negative loading matrix for spatial domain 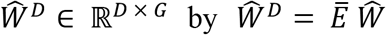 . The column-normalization and gene ranking are performed similarly to the procedure described in the section, “Ranking and identification of genes associated with embedding dimensions”.

### High-resolution gene expression recovery

For the *g* -th gene, SpaHDmap recovers its high-resolution normalized expression 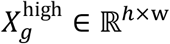by computing a weighted sum of the learned embeddings *F*^high^,

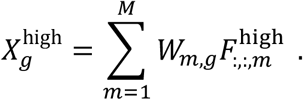

Note that 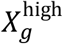quantifies the expression of the *g*-th gene for every pixel in the tissue, and thus essentially imputes the sparse expression quantification given by the original ST expression data.

### Joint analysis of multi-sample ST data

In the case of multiple ST samples, the SpaHDmap framework remains similar as before, with the addition of learnable batch parameters to account for potential batch effects across different samples. Specifically, let *Ω*_*b*_ be the sequenced tissue of the *b*-th sample (*b* = 1,2, … , *B*). Similar to Equation (2), SpaHDmap reconstructs the normalized gene expression 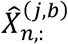of the *n*-th spot in the *j*-th tissue patch from the *b*-th sample by employing a non-negative linear combination component alongside an additional batch effect term as follows:

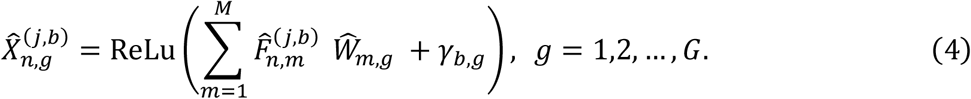

Here, 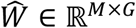 is a non-negative loading matrix to be learned, 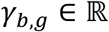denotes the learnable mean batch effect parameter of batch *b* on gene *g*. For identifiability, we set the first batch as the reference batch, i.e., γ_1,*g*_ = 0. Further, the spot-wise low-resolution factor matrix *F*^(b)^ of batch *b*, derived from Equation (1), is replaced by

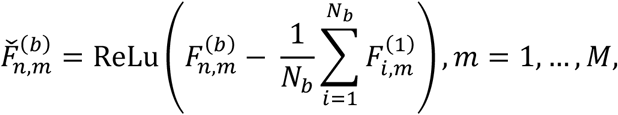

where *N*_*b*_ is the number of spots in the reference batch.

For samples from *C* biologically different experimental conditions (e.g., Palbociclib-treated and untreated conditions in the SHH medulloblastoma dataset), where there are *B*_*c*_ batches for condition *c*, we replace *γ*_*b*,*g*_ in Equation (4) with *γ*_*c*,*b*,*g*_ ∈ ℝ (*c* = 1, … , *C*; *b* = 1, … , *B*_*c*_) as follows:

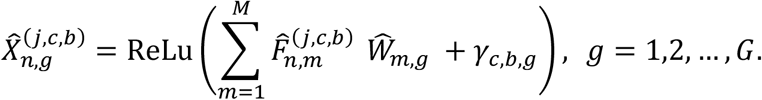

For identifiability, we set the first batch in each condition as the reference batch, i.e., γ_*c*,1,*g*_ = 0. All parameters are trained using the Adam optimizer to minimize the overall reconstruction loss (3).

### Generation and analysis of simulated datasets

Two methods are used to generate simulation datasets. The first method utilizes an H&E image of a mouse cerebellum and generates low-resolution ST expression data using the zero-inflated Poisson distributions, where the distribution parameters are dependent on the given H&E image. The second method uses a real single-cell-resolution ST dataset sequenced by the 10X Xenium to generate low-resolution ST expression data.

### Simulation data generation using an H&E image of mouse cerebellum

We utilize an H&E image of a mouse posterior brain sagittal section (4000×4000 pixels) sequenced by 10X Visium(https://www.10xgenomics.com/resources/datasets/mouse-brain-serial-section-1-sagittal-posterior-1-standard-1-1-0) to simulate ST expression data. The image is first segmented into four regions: fiber tracts, granular layer of the cerebellum, molecular layer of the cerebellum, and background area. We introduce *M* = 3 true pixel-wise embedding dimensions, denoted as *F*^*^ ∈ ℝ^*h* × *w*× *M*^, where *h* = 4000 and *w* = 4000 are the height and width of the image, respectively. Each dimension corresponds to and is only enriched in one specific segmented region, excluding the background area.

We generate *N* spots with radius *r* using a 10X Visium-style meshgrid. Denote the pixel set located in the *n*-th spot as *S*_*n*_ (*n* = 1, … , *N*). Using the pixel-wise embedding *F*^*^, we induce the spot-level embedding dimension intensity 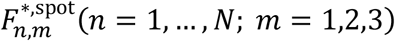 as follows,

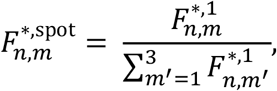

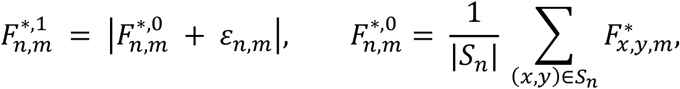

where ε_*n*,*m*_ ∼𝒩(0, 0.05^2^) is a Gaussian random variable that controls spot-specific variation of the embedding intensity. Then, we generate the loading matrix 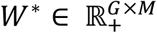 as follows. We divide the *G* genes into 3 gene sets, and assign each gene set to one embedding as its associated gene set. The loading 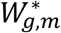 is generated from a Gamma distribution with a scale parameter of 0.05. The shape parameter of the Gamma distribution is set to 15 if the gene is associated with the dimension, and 5 otherwise.

Given the generated spot-level embedding *F*^*,spot^ and the loading matrix *W*^*^, we first generate the true underlying log-transformed gene expression *X*_*n*,*g*_ for spot *n* and gene *g* as 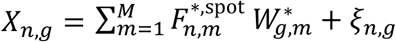, where *ξ*_*ng*_ is independently generated from 𝒩(0, *σ*^2^). Then, we generate the observed count expression 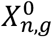 for spot *n* and gene *g* from a zero-inflated Poisson distribution. The zero-inflation parameter of the zero-inflated Poisson distribution is *π* and the mean parameter is *λ* = *l*_*n*_ exp *X*_*n*,*g*_ − 1, where *l*_*n*_ represents the spot-specific library size and is generated using log(*l*_*n*_)∼𝒩(10, 0.05^2^) . After generating the count expression data, we also randomly exclude a proportion *ρ* of the spots to simulate different spot densities.

In the simulation, we set the number of genes as *G* = 3000. We vary the spot exclusion percentage *ρ*, the spot radius *r*, the expression variance *σ*, and the zero-inflation parameter *π* to generate different simulation scenarios. For each scenario, we generate 30 replicates.

### Simulation based on a real single-cell-level ST dataset

We use a single-cell-level Xenium ST dataset from a human breast carcinoma tissue^3^ to generate simulated spot-level ST expression profiles. This Xenium ST dataset contained an expression profile for 313 genes across 167,780 sequenced cells, and an H&E image with 5478 × 7526 pixels. To generate the simulated spot-level ST expression data, we first generate spots *S*_*n*_ (*n* = 1, … , *N*) with a radius *r* using a 10X Visium-style meshgrid and sum the expression counts of cells within each spot as the spot-level expression. Then, a proportion *ρ* of the generated spots are excluded along with their spatial coordinates and gene expression.

Considering that the number of annotated cell types for this Xenium dataset is 19, we set the number of embedding dimensions as *M* = 20 in this simulation. The underlying true single-cell-level embedding is derived from applying NMF to the original Xenium ST dataset. We vary the spot exclusion proportion *ρ* and the spot radius *r* to generate different simulation scenarios, generating 30 replicates for each scenario.

### Evaluation criteria of the embedding learning methods

We use the MAE between the true embedding intensity *F*^*^ and the inferred embedding intensity 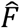 to evaluate each method’s performance on embedding learning. The MAE for one simulated dataset is defined as

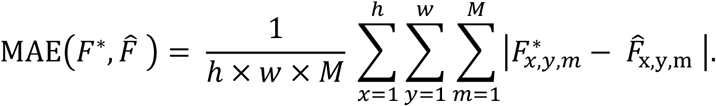

For methods that only give spot-level embeddings, we extend their spot-level embeddings to entire-tissue-wide embeddings using Voronoi tessellations and rasterization before calculating their MAEs. To assess the accuracy of the spatial domain assignment in simulation data generated from mouse cerebellum, we cluster the pixels into three regions by applying the K-means algorithm to the inferred embedding of each method, and then compare the inferred region label with the three true regions using the ARI metric.

### Comparison with available methods

We compare SpaHDmap with various interpretable embedding learning and spatial domain detection methods. The interpretable embedding learning methods include NMF (via the *decomposition*.*NMF* function in the Python *scikit-learn* package, v1.5.1), NSFH, SpiceMix, Giotto (v4.0.0) and TESLA (v1.2.4)+NMF. The spatial domain detection methods include iStar, stLearn (v0.4.12) and SpaGCN (v1.2.7). NMF, NSFH, SpiceMix and Giotto only require ST expression data, while iStar, stLearn, TESLA and SpaGCN require both ST expression data and histology image. The available methods that utilize histology images all require RGB images as input. Thus, for datasets with H&E images, we directly provide the H&E images to these methods, while for datasets with IHC images, we first convert the IHC images to RGB images and provide the obtained RGB images.

In terms of embedding learning, for TESLA+NMF, we first use TESLA to enhance the spatial resolution of gene expression based on the histology image and spatial expression profile. The default meta-pixel size of 50 × 50 pixels is used for TESLA. Then, we learn an embedding using the standard NMF based on the resolution-enhanced expression. For other embedding learning methods, their default parameters are used.

In terms of the spatial domain detection, for the embedding learning methods, we first apply these methods to obtain the embeddings. Then, for the spot-level embedding methods, we apply the Louvain algorithm^59^ to their embeddings for spot clustering. For the high-resolution embedding methods including SpaHDmap and TESLA+NMF, we first apply the Louvain algorithm to cluster the spots and then apply the K-means algorithm to cluster the meta-pixels with the clustering results given by Louvain as the initial values. Across all embedding methods, we use the same resolution parameter for the Louvain algorithm and the same cluster number of the K-means algorithm for clustering. For the spatial domain detection methods, we employ their default parameters and specify the same number of clusters as that of the embedding methods for spatial domain identification.

### Sample collection and library preparation of colorectal cancer

CRC-01 was sampled from the abdominal wall of a metastatic colorectal cancer patient, while CRC-02 was sampled from the ovarian metastasis site of another colorectal cancer patient. Formalin-fixed paraffin-embedded (FFPE) samples of CRC-01 and CRC-02 were sectioned into 5 μm slices by Leica HistoCore BIOCUT and were affixed to adhesive slides. Subsequently, the FFPE sections underwent spatial transcriptomics profiling using Visium CytAssist Spatial Gene Expression Reagent Kits (10X Genomics, USA) following the manufacturer’s protocols and recommended third party reagents. Visium spatial libraries were sequenced in paired-end with Illumina NovaSeq 6000 according to manufacturer instructions.

CRC-03 was sampled from a primary colorectal cancer patient. Fresh tissue sample of CRC-03 was obtained and embedded in Tissue-Teck OCT within 30 minutes of resection, with extra fluid being wiped out. The sample was then frozen in dry ice and transferred to a -80° C freezer for storage. Freezing and OCT-embedding of the tissue sample were sectioned to 10μm slices by Leica CM1950 cryostat, and immediately attached to Stereo-seq chips. The target tissue section was fixed with formaldehyde, and stained using nucleic acid dye (Thermo fisher, Q10212). The section was permeabilized for RNA release at the selected time to capture the mRNA molecules from the target tissue cells on the microarray for cDNA synthesis. The indexed cDNA libraries were constructed and sequenced in paired-end with MGI DNBSEQ-T7 sequencer according to manufacturer instructions.

### Generation of the pseudo-image for the colorectal cancer Stereo-seq dataset

The Stereo-seq dataset CRC-03 lacks paired histology images. To apply SpaHDmap and other image-depended methods, we first generated a high-resolution pseudo-image based on the subcellular expression profile of Stereo-seq data. Specifically, we first extracted the high-resolution bin-level expression counts using the *io*.*read_gef* function from the Python *Stereopy* package with a bin size of This bin size is significantly smaller than the bin size commonly used for Stereo-seq data analysis (i.e., bin size = 100). Next, the bin-level expression counts were log-transformed and normalized using the total sum count as the library size. Principal Component Analysis (PCA) was then performed on the bin-level normalized expression. Finally, we summarized the top 30 PCs into three UMAP components and visualized the three-dimensional components using red, green, and blue (RGB) colors through an RGB plot. This process resulted in a pseudo-image that captures the high-resolution spatial structure of the tissue (Fig. S19a).

### Analysis of colorectal cancer samples

Cell type proportion deconvolution was performed using RCTD^44^ in the R package *spacexr* (v2.2.1). The reference single-cell RNA-seq data for RCTD was downloaded from the National Center for Biotechnology Information (NCBI) Gene Expression Omnibus (GEO) under accession code GSE183202^60^.

To characterize different biological activities, we computed the mean expression of gene lists provided in Supplementary Table 3 using the *AddModuleScore* function in the R package *Seurat* (v4.0.0). Given the embeddings from SpaHDmap, we calculated the mean ranks of the given gene lists in each high-resolution cluster. The gene lists include genes related to CMS subtypes^42, 43^, functional states of cancer cells from CancerSEA^61^ and cell-type markers^47^ (Supplementary Table 3). Copy number variation detection was carried out using the R package *infercnv* (v1.20.0, https://github.com/broadinstitute/inferCNV).

For survival analysis, we downloaded TCGA^46^ expression and survival data using the R package *cBioPortalData*^62-64^. For each gene, we standardized its expression to have a mean of 0 and standard deviation of 1 across samples. Given a list of genes, we first calculated their mean expression for each patient and the upper and lower quantiles of its mean expressions across the patients. Patients with a mean expression larger than the upper quantile were assigned to the high expression group, while patients with a mean expression smaller than the lower quartile were assigned to the low expression group. Kaplan-Meier curves and log rank tests were carried out using the *survfit* and *ggsurvplot* functions in the R packages *survival* and *survminer*, respectively.

CMS classification was performed using the R package *CMSclassifier* (v1.0.0)^42^. Briefly, the raw count matrix of RNA-seq was transformed by variance stabilizing transformation^65^ (*SCTransform*). Then, gene identifiers were converted from gene symbols to Entrez gene ids using the R package *org*.*Hs*.*eg*.*db* (genome-wide annotation for Human, v3.19.1). The processed matrix was used as input for the *classifyCMS* function with the option method “SSP”.

## Data description

The first simulation dataset was generated based on the H&E image of a mouse brain dataset MPBS-01 from the 10X Visium platform, while the second simulation dataset was generated from a human breast cancer ST dataset with an H&E image sequenced by 10X Xenium. The mouse brain datasets MBC-01, MBC-02, MBC-03, MPBS-01 and MPBS-02 were sequenced by 10X Visium, with IHC images for MBC-01 and MBC-02, and H&E images for the other mouse brain datasets. The four SHH medulloblastoma PDOX datasets were sequenced by 10X Visium and accompanied with H&E images, with their tumor/normal/interface annotations from the original paper^39^. The human colorectal cancer samples CRC-01 and CRC-02 were sequenced by 10X Visium CytAssist. Sample CRC-03 was sequenced by Stereo-seq technology with a nearby slice stained by H&E image. See Supplementary Table 1 for details.

The region annotations for mouse brain coronal and sagittal sections were downloaded from the aligned slices (the 76th coronal section and the 9th sagittal section) in the Allen Reference Atlas – Mouse Brain^66^ (atlas.brain-map.org). The mouse brain region marker genes were sourced from the Allen Mouse Brain Atlas^31^ (https://mouse.brain-map.org/). The marker genes of major cell types in the adult mouse leptomeninges were collected from Pietila et al.^36^. The marker genes of mouse microglial cells, astrocytes, and endothelial cells were downloaded from the CellMarker 2.0 database (http://117.50.127.228/CellMarker/index.html). The gene signatures of SHH-A, SHH-B, SHH-C, E2F target genes, and neuronal differentiation were collected from Vo et al.^39^. Reference scRNA-seq of metastasized colorectal cancer samples were downloaded from the NCBI GEO database under accession code GSE183916.

## Data availability

The ST datasets in the paper are publicly available at:

1. 10X Visium mouse brain coronal section data. MBC-01: https://www.10xgenomics.com/resources/datasets/adult-mouse-brain-section-2-coronal-stains-dapi-anti-gfap-anti-neu-n-1-standard-1-1-0 MBC-02: https://www.10xgenomics.com/resources/datasets/adult-mouse-brain-section-1-coronal-stains-dapi-anti-neu-n-1-standard-1-1-0 MBC-03: https://www.10xgenomics.com/resources/datasets/mouse-brain-section-coronal-1-standard-1-0-0
2. 10X Visium mouse posterior brain sagittal section data. MPBS-01: https://www.10xgenomics.com/resources/datasets/mouse-brain-serial-section-1-sagittal-posterior-1-standard-1-1-0 MPBS-02: https://www.10xgenomics.com/resources/datasets/mouse-brain-serial-section-2-sagittal-posterior-1-standard-1-1-0
3. 10X Xenium human breast cancer data, replicate 1. https://www.10xgenomics.com/products/xenium-in-situ/preview-dataset-human-breast
4. 10X Visium sonic hedgehog medulloblastoma data. https://genomemedicine.biomedcentral.com/articles/10.1186/s13073-023-01185-4#availability-of-data-and-materials

The raw and processed data of colorectal cancer samples will be available at the time of publication. Details of the datasets analyzed in this paper are described in Supplementary Table 1.

## Code availability

An open-source implementation of the SpaHDmap algorithm can be accessed at https://github.com/XiDsLab/SpaHDmap. A detailed documentation, including tutorials demonstrating the use of SpaHDmap on several example datasets, is available at https://spahdmap.readthedocs.io/en/latest/. Codes to reproduce the analyses in this manuscript are available at Zenodo: https://doi.org/10.5281/zenodo.13690848.

## Acknowledgments

This work was supported by the National Natural Science Foundation of China (12425110, 12371286 and 11971039 to R.X.), the National Key R&D Program of China (2020YFE0204200 to R.X.), and Science and Technology Project of Tianjin Binhai New Area Health Commission (2022BWKY016 to Y.Z.). Part of the analysis was performed on the high-performance computing platform of the Center for Life Sciences (Peking University). We thank Quan Nguyen and Laura A. Genovesi for sharing the 10X Visium mouse brain sonic hedgehog (SHH) medulloblastoma data. We also thank Yuanyuan He and Cheng Li for their helpful discussions.

## Author contributions

R.X. conceived the study. R.X. and J.J.X. supervised the study. J.T. formulated the method. J.T. and K.Q. developed the software. J.T. and Z.C. performed bioinformatic analyses and prepared figures and tables in the manuscript. S.H. helped develop biological interpretations. S.Y., B.Y. and H.M. conducted pathological annotations. Y.Z. prepared the samples. Y.H. performed the spatial transcriptomics sequencing. J.T., Z.C., X.H. and R.X. wrote the manuscript.

## Ethics declarations

All human tissue samples were obtained from Tianjin Medical University Cancer Institute & Hospital. Before surgery at the center, all patients provided written informed consent to allow any excess tissue to be used for research studies. The study was approved by the Ethics Committee (ID: bc2022189).

